# Correcting axial-motion artifacts in two-photon calcium imaging with suite2p-in-depth

**DOI:** 10.64898/2026.09.25.754364

**Authors:** Liad J. Baruchin, Maria Cozan, Andrew Mikhniak, Carsen Stringer, Sylvia Schröder

## Abstract

Two-photon calcium imaging enables measurements of neural activity from individual neurons and their subcellular processes in awake animals. Axial brain motion can change measured fluorescence independently of neural activity and move small structures out of the imaging plane, confounding two-photon calcium measurements in behaving animals. We developed two complementary post-acquisition methods, selected according to whether the imaged structure remains detectable across the encountered depth range. When it does, z-stack correction estimates imaging depth from a separately acquired reference z-stack and uses an axial fluorescence profile specific to each neural structure to rescale fluorescence to a common reference depth. When a structure can leave a single imaging plane, volume-based z-registration reconstructs a common-depth movie from repeated acquisitions of closely spaced planes. In awake mice, both methods tracked experimentally imposed axial displacements within the sampled range, with depth-estimation root mean squared errors (RMSEs) below 1 *µ*m for z-stack correction and 0.69–1.03 *µ*m for volume-based z-registration. Correction changed fluorescence substantially for a subset of somata, with larger effects in axonal boutons. Both methods are implemented in suite2p-in-depth, a Python package that integrates with Suite2p and provides diagnostics for depth estimation, correction quality and data reliability. Together, these methods enable axial-motion artifacts to be assessed and corrected within established Suite2p workflows.

## Introduction

Understanding how neural circuits support behavior requires measurements of activity at both cellular and subcellular scales. Two-photon imaging with genetically encoded calcium indicators provides access to signals from neuronal somata, dendrites, axons and presynaptic boutons in behaving animals (Broussard and Petreanu, 2021; Stringer and Pachitariu, 2019). Interpreting these measurements requires separating activity-dependent fluorescence changes from artifacts caused by movement of brain tissue relative to the microscope’s focal plane. Lateral movement can be compensated by aligning images within the imaging plane, but alignment does not address movement perpendicular to it. Such axial motion changes the fraction of a fluorescent structure captured in the optical section and therefore alters its measured signal independently of neural activity. Axons and boutons are especially vulnerable because they can leave the imaging plane entirely (Broussard and Petreanu, 2021; Ryan et al., 2020).

Axial motion occurs even in head-fixed animals and has been documented during locomotion and licking (Chen et al., 2013; Kong et al., 2016; Ryan et al., 2020). Because it can covary with behavior, depth-dependent fluorescence changes may mimic behavioral modulation or obscure genuine activity changes. Correcting axial-motion artifacts has been shown to change the apparent relationship between synaptic calcium signals and running speed (Ryan et al., 2020). Reliable interpretation therefore requires assessing both axial movement and its effects on the fluorescence of individual structures.

Axial motion can be addressed during acquisition or corrected offline. Online stabilization adjusts the sampled position using behavioral signals or direct optical tracking (Chen et al., 2013; Gonzalez et al., 2026; Griffiths et al., 2020; Stringer et al., 2026). These approaches require focus-control hardware and software, and a tracking error can cause irrecoverable signal loss if the acquisition no longer samples the structure of interest. Offline approaches instead retain the original measurements and have used reference z-stacks, multiple focal planes or repeated volumes to estimate displacement and correct or register the data after acquisition (Andermann et al., 2010; Flores-Valle and Seelig, 2022; Ryan et al., 2020).

Here, we present suite2p-in-depth, a Suite2p-compatible Python package implementing two complementary offline approaches for axial-motion correction: z-stack correction of ROI fluorescence and volume-based z-registration of multiplane recordings. We validate both methods against experimentally imposed displacements and assess their effects in recordings of neuronal somata and retinal axonal boutons in the superior colliculus (SC) of awake mice.

## Results

We developed two complementary approaches to correct the effects of axial motion, chosen according to whether the structure of interest remains detectable over the range of depths encountered during a recording (**Figure 1**). If a structure remains detectable throughout this range, axial motion changes the fraction sampled by the imaging plane and therefore modulates the fluorescence measured within its ROI mask, as often occurs for neuronal somata (**Figure 1**a,b). If axial movement carries a structure partly or completely out of the imaging plane, its measured fluorescence can instead be strongly attenuated or lost, as often occurs for axonal boutons (**Figure 1**c,d). The appropriate correction strategy therefore depends on the structure’s axial fluorescence profile relative to the motion. Z-stack correction addresses the first case by estimating imaging depth from a separately acquired reference z-stack and using an ROI-specific axial fluorescence profile to rescale fluorescence to a common reference depth (**Figure 1**e,g). Volume-based z-registration addresses the second case by repeatedly sampling closely spaced planes and reconstructing a depth-stabilized movie from frames that best represent a common reference depth (**Figure 1**f,h).

**Figure 1.**
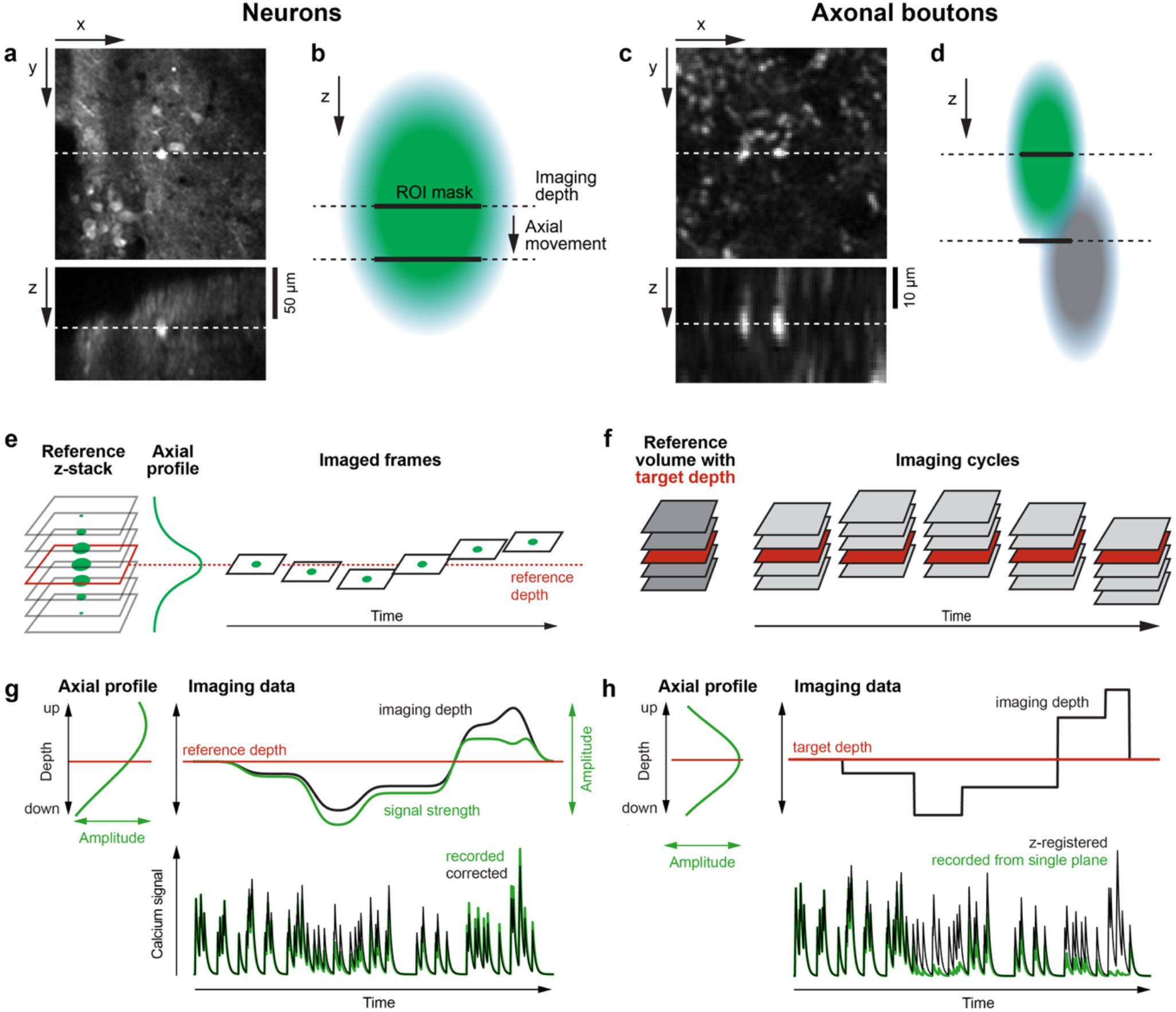
Axial motion and the two correction strategies. **a**. Example x-y and x-z views of GCaMP8m-expressing neurons in the mouse superior colliculus (SC). **b**. Schematic showing how axial displacement changes fluorescence measured within a fixed neuronal ROI mask. **c**,**d**. Corresponding example and schematic for SyGCaMP8m-labeled retinal axonal boutons, which can move out of a single imaging plane. **e**. Z-stack correction estimates imaging depth by matching functional frames to a reference z-stack and derives an ROI-specific axial fluorescence profile. **f**. Volume-based z-registration matches repeated multiplane imaging cycles to a reference volume and selects frames corresponding to a common target depth. **g**. Z-stack correction uses the estimated imaging depth and axial fluorescence profile to correct depth-dependent fluctuations in the recorded fluorescence. **h**. Volume-based z-registration reconstructs fluorescence at the target depth by selecting the best-matching plane from each imaging cycle.

The two approaches enter the Suite2p workflow at different stages: z-stack correction operates on data already processed with Suite2p, whereas volume-based z-registration reconstructs a depth-stabilized movie that is subsequently processed with Suite2p (Stringer et al., 2026). Both workflows support manual ROI curation through the Suite2p GUI.

### A. Z-stack correction for structures that remain detectable across depth

Z-stack correction is applied after standard Suite2p processing of the functional recording. The recording is first processed with Suite2p to laterally register frames, generate a reference image, detect ROIs and extract ROI and neuropil fluorescence traces. These outputs, together with a separately acquired reference z-stack, are then passed to the z-stack correction workflow. The algorithm estimates imaging depth from the reference z-stack and uses an ROI-specific axial fluorescence profile to correct ROI and neuropil fluorescence to a common reference depth before neuropil correction and calculation of ΔF/F (**Figure 2**).

**Figure 2.**
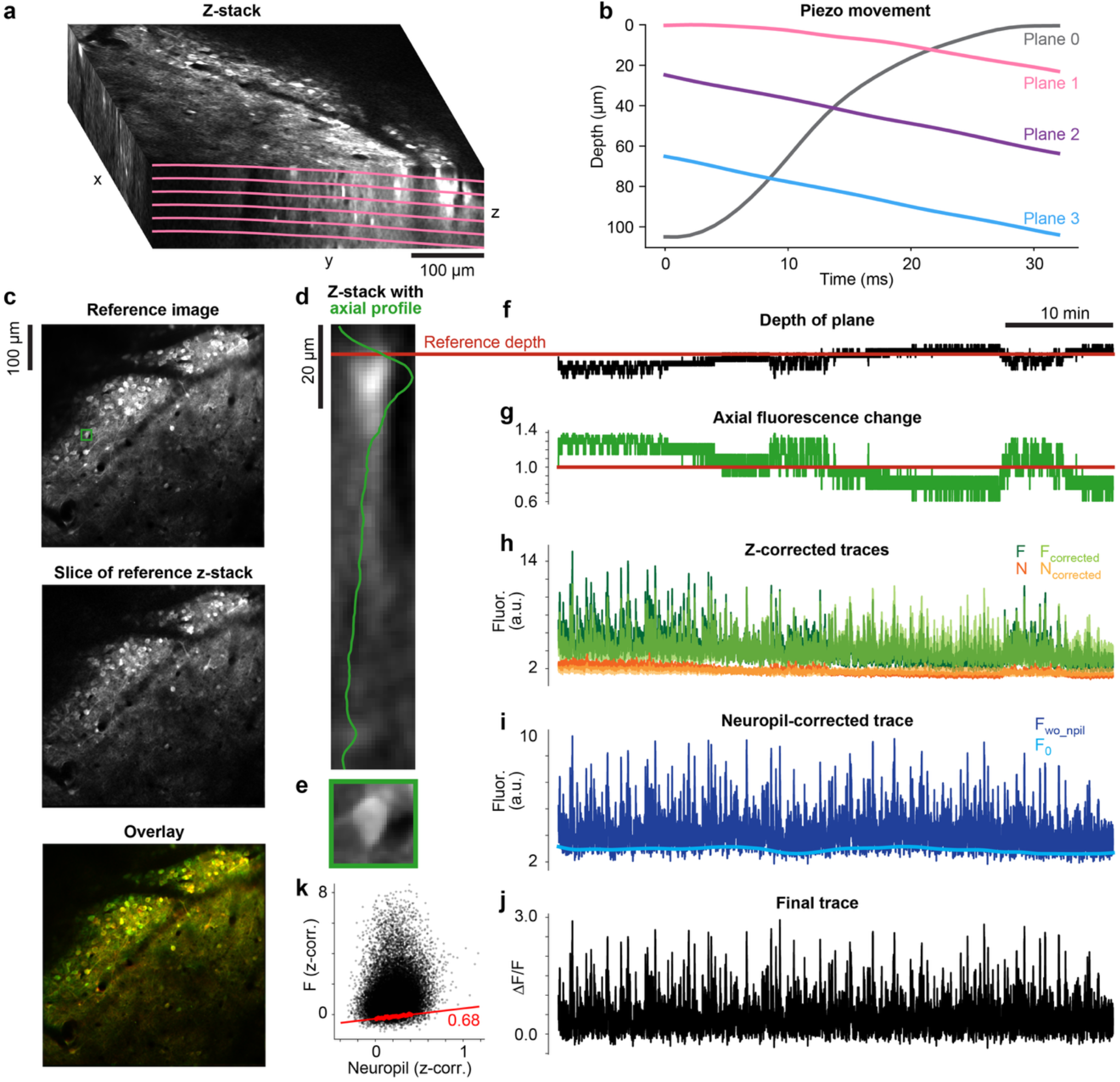
Z-stack correction workflow. **a**. Reference z-stack and depth profile of an example functional imaging plane (pink), obtained from the piezo trajectory in **b. b**. Piezo position during acquisition of the functional imaging planes; plane 0 is a flyback plane. **c**. Suite2p reference image (top), corresponding slice of the resliced z-stack (center), and their overlay after lateral alignment (bottom). The boxed ROI is used in **d–k. d**,**e**. Axial view (**d**) and top view (**e**) of the example ROI. The green line in **d** shows its axial fluorescence profile and the red line the reference depth. **f**. Estimated imaging depth over time. **g**. Depth-dependent correction factor derived from the ROI’s axial profile. **h**. ROI and neuropil fluorescence before and after axial correction. **i–k**. Subsequent neuropil correction (**i**) and calculation of ΔF/F (**j**); **k** shows the fit used to estimate the ROI-specific neuropil coefficient.

#### Z-stack correction workflow

1. *Construct a reference z-stack matching the functional imaging plane (**Figure 2**a-c)* Repeated images acquired at each z-stack depth are laterally aligned and averaged, and the resulting planes are aligned across depth to form a registered reference volume. If the functional imaging plane is not parallel to the z-stack planes, as can occur during continuous piezo scanning, the volume is resliced to reproduce the geometry of the functional plane. The slice that best matches the corresponding Suite2p reference image is then identified and used to laterally align the resliced z-stack to the functional data.
2. *Estimate imaging depth over time (**Figure 2**f)* Each laterally registered functional frame is compared with the slices of the aligned reference z-stack and assigned the depth of its best-matching slice. This yields an estimated imaging-depth trace for each functional plane. The median estimated depth across the recording is used as the reference depth to which fluorescence is subsequently corrected.
3. *Derive ROI-specific axial fluorescence profiles (**Figure 2**d,e,g)* For each ROI, its mask is applied to every slice of the aligned reference z-stack to measure fluorescence as a function of depth. The resulting axial profile provides an ROI-specific calibration of the fluorescence changes expected from axial displacement.
4. *Correct fluorescence to the reference depth (**Figure 2**h)* At each time point, the ROI fluorescence is divided by the axial-profile amplitude at the estimated depth relative to its amplitude at the reference depth. The same correction factor is applied to the corresponding neuropil signal. This rescales both signals to the fluorescence expected at the reference depth. Measurements are retained only when the estimated depth lies within the reliable region of the ROI’s axial profile; measurements outside this region are set to NaN rather than extrapolated (see Methods).
5. *Perform neuropil correction and calculate ΔF/F (**Figure 2**i-k)* After axial correction, slow baseline components are estimated for the ROI and neuropil traces. An ROI-specific neuropil coefficient is then estimated from their relationship and used to subtract neuropil contamination from the ROI signal. The resulting fluorescence trace is converted to ΔF/F (see Methods).

#### Performance and validation

We evaluated z-stack correction using two-photon imaging of GCaMP8m-expressing superior colliculus (SC) neurons in awake, head-fixed mice able to run voluntarily on a treadmill. All but one mouse expressed tdTomato in inhibitory neurons, providing a stable, activity-independent signal for lateral registration and axial matching; in the remaining mouse, GCaMP fluorescence was used instead. A piezoelectric actuator sampled three functional imaging planes at 7.5 Hz per plane. Separate recordings were used for ground-truth validation and for quantifying the effects of correction on fluorescence measurements.

In the two ground-truth recordings, the algorithm’s depth estimates closely tracked axial displacements manually imposed with the piezoelectric actuator (**Figure 3**a). The imposed displacements were superimposed on the sawtooth trajectory used for multiplane acquisition. Assuming that brain motion was negligible relative to the imposed displacement, the actuator trajectory provided the ground-truth imaging depth. To account for the arbitrary depth zero point, we fitted a single constant offset shared across the three planes; no scaling or time-varying parameters were fitted. The RMSEs between estimated and ground-truth depths ranged from 0.39 to 0.92 *µ*m across the six plane-recording combinations.

**Figure 3.**
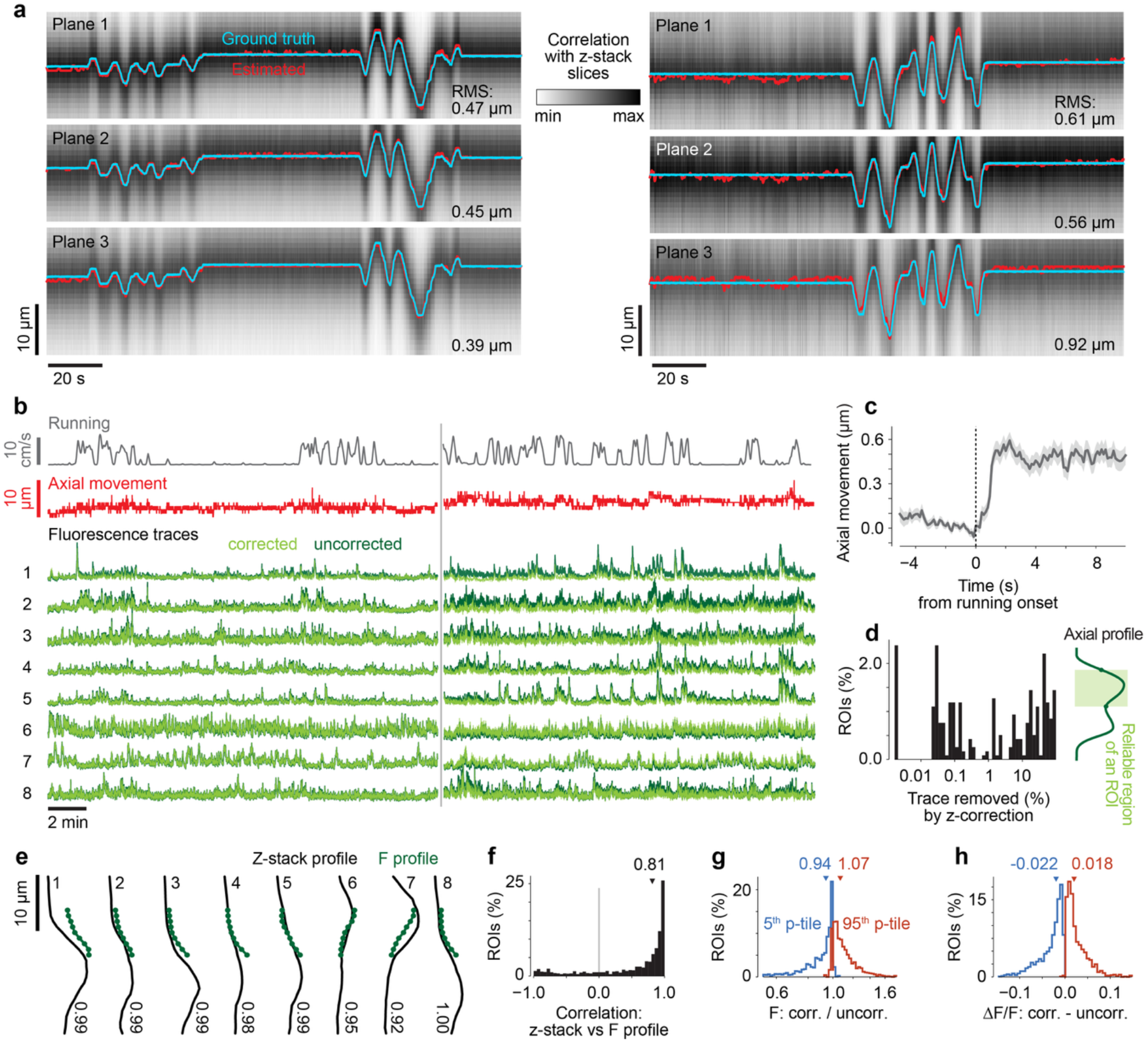
Validation and effects of z-stack correction. **a**. Estimated (red) and experimentally imposed ground-truth (cyan) axial displacement in two recordings. Rows show the three functional imaging planes; grayscale indicates the correlation of each frame with slices of the corresponding reference z-stack. RMSEs are indicated. **b**. Running speed, estimated axial displacement, and uncorrected (dark green) and z-stack-corrected (light green) fluorescence from eight example ROIs in one functional recording. The vertical gray line separates two drifting-grating paradigms. **c**. Estimated axial displacement aligned to 140 running onsets from the recording in **b** (mean ± SEM). **d**. Percentage of time points excluded because the estimated depth fell outside the reliable region of the ROI’s axial profile or reached a z-stack boundary. Only ROIs with at least one excluded time point are shown (306/1,179 ROIs); schematic at right illustrates the reliable axial-profile region. **e**. Axial fluorescence profiles for the eight ROIs in **b**, obtained from the reference z-stack (black) and from functional fluorescence binned by estimated depth (green). Numbers indicate Pearson correlation coefficients. **f**. Distribution of profile correlations across 1,179 ROIs from five recordings; triangle indicates the median. **g**. Across-ROI distributions of the 5th (blue) and 95th (orange) percentiles, calculated over time, of the ratio between corrected and uncorrected fluorescence. **h**. Corresponding distributions for the difference between corrected and uncorrected ΔF/F.

In five additional functional recordings from five mice, estimated axial position varied both in association with running and independently of it (**Figure 3**b,c). Running onset was often accompanied by an axial displacement whose direction was consistent within a recording but varied across recordings, with upward movement in the example shown and downward movement in others. Axial position also exhibited slow drift and shifts between consecutive recording blocks, which were typically separated by a few minutes.

We next quantified how often the correction identified measurements as unreliable (**Figure 3**d). Across 1,179 ROIs, 306 (26%) had at least one time point excluded because the estimated depth lay outside the ROI’s reliable axial-profile region or reached a boundary of the reference z-stack. 9% of all ROIs had at least 10% of their time points excluded, whereas the remaining 74% could be corrected throughout the recording.

Axial fluorescence profiles derived from the reference z-stack showed substantial agreement with profiles reconstructed from the functional imaging data (**Figure 3**e,f). This comparison tests a central assumption of z-stack correction: that the relationship between ROI fluorescence and imaging depth measured in the reference z-stack remains applicable during functional imaging. To reconstruct the functional-imaging profile for each ROI, we binned its uncorrected fluorescence according to estimated imaging depth and calculated the median within each depth bin. The median Pearson correlation between the paired profiles was 0.81 (interquartile range, 0.26–0.95; **Figure 3**f). This agreement was observed even though reference z-stacks were sometimes acquired up to two hours before or after the corresponding functional-imaging blocks and activity-dependent fluorescence differed between the acquisitions.

Z-stack correction produced modest changes in fluorescence and ΔF/F for most ROIs but substantially larger changes in a subset. To quantify the effect on fluorescence, we calculated the ratio of corrected to uncorrected fluorescence at each valid time point and summarized its distribution within each ROI by its 5th and 95th percentiles (**Figure 3**g). Across ROIs, the median 5th- and 95th-percentile ratios were 0.94 and 1.07, corresponding to changes of −6% and +7%, respectively. However, 21% of ROIs had a 5th-percentile ratio below 0.8, meaning that correction reduced fluorescence by more than 20% for at least 5% of valid time points. Conversely, 17% had a 95th-percentile ratio above 1.2, indicating increases of more than 20% for at least 5% of valid time points.

We similarly quantified the difference between corrected and uncorrected ΔF/F after neuropil correction and baseline normalization (**Figure 3**h). The median 5th- and 95th-percentile differences were −0.02 and +0.02, respectively; 22% of ROIs had a 5th-percentile difference below −0.05 and 15% had a 95th-percentile difference above +0.05.

#### Data requirements and practical considerations

Z-stack correction requires two matched inputs: a functional recording processed with Suite2p and a reference z-stack that overlaps the same field of view and can be geometrically aligned to the functional imaging plane.

##### Reference z-stack acquisition

Acquire a densely sampled z-stack over the same field of view as the functional recording. The stack should extend beyond the expected range of axial motion, with sufficient margin above and below the functional plane, and its axial sampling interval should be small enough to resolve changes in image similarity and ROI fluorescence across depth.

The z-stack may be acquired using a motorized objective or stage, a piezo objective drive, or another remote-focusing system. Imaging parameters that affect image geometry or axial fluorescence profiles—including objective, excitation wavelength, zoom, scan rotation, pixel size and field of view—should be kept as consistent as possible between the z-stack and functional recording. The stack should ideally be acquired close in time to the functional recording and over a sufficiently short interval to minimize tissue movement and changes in imaging conditions.

##### Matching the imaging-plane geometry to the z-stack

If the functional imaging plane is parallel to the z-stack planes, no additional geometric information is required. During continuous axial scanning, however, the objective can move appreciably within a frame, producing a functional imaging plane that is not parallel to the z-stack planes. In this case, the reference volume must be resliced to reproduce the functional-plane geometry. This requires synchronized measurements of axial-actuator position and functional frame timing; in our setup, these were provided by the piezo-position trace and frame-clock signal.

#### Assumptions, limitations, and quality control

The validity of the z-stack correction depends on reliable estimation of imaging depth and on the reference z-stack providing an appropriate calibration of depth-dependent fluorescence. These requirements should be evaluated at the level of both the recording and the individual ROI.

1. Functional frames must support reliable registration and depth estimation Functional frames must contain sufficiently stable image structure to permit reliable lateral registration to the corresponding Suite2p reference image and subsequent matching to the reference z-stack. If changes in image appearance prevent reliable registration or produce ambiguous depth matches, both the estimated depth and resulting fluorescence correction become unreliable.
2. The axial fluorescence profile is treated as a multiplicative gain function The correction assumes that imaging depth acts as a multiplicative gain on the ROI fluorescence, described by the ROI’s axial profile (Ryan et al., 2020). Under this model, changes in neural activity scale fluorescence without substantially changing the shape of the axial profile. Violations of this assumption can bias the corrected trace.
3. Correction is restricted to a reliable region of each ROI’s axial profile The axial fluorescence profile provides a reliable correction only over a limited range of depths around the reference depth. At more distant depths, fluorescence may become too weak for stable correction, or the two-dimensional ROI mask may overlap a different structure across depth. The algorithm therefore defines an ROI-specific reliable region around the reference depth and applies correction only when the estimated imaging depth lies within this region. Measurements outside the reliable region are set to NaN rather than extrapolated (see Methods).
4. The reference z-stack must remain a valid calibration The relationship between imaging depth and measured fluorescence captured by the reference z-stack must remain applicable during functional recording. Tissue movement during z-stack acquisition, inaccurate alignment, changes in imaging geometry, or changes in the structure’s appearance can invalidate this calibration. Subsequent signal processing cannot compensate for an incorrectly aligned or otherwise unrepresentative axial profile.
5. Quality-control checks Before interpreting corrected traces, users should inspect the alignment between the reference z-stack and the functional reference image (**Figure 2**c), the frame-to-stack correlations and estimated depth traces (**Figure 3**a,b), the frequency of matches at the z-stack boundaries, and the shape and reliable region of each ROI’s axial profile (**Figure 2**d). The proportion of each trace set to NaN and the differences between corrected and uncorrected fluorescence should also be quantified. Poor alignment, persistent boundary matches, ambiguous axial profiles, extensive data loss, or correction-induced discontinuities or noise indicate that the correction may be unreliable for that recording or ROI. The suite2p-in-depth package automatically generates diagnostic outputs for these checks, including plots similar to those shown in **Figure 2** and **Figure 3**. Guidance for interpreting these outputs is provided in the GitHub repository.

#### B. Volume-based z-registration for structures that can leave a single imaging plane

Volume-based z-registration is used when axial motion can move structures partly or completely out of a single imaging plane, as often occurs for axons, boutons and dendrites. The method is applied before ROI detection and requires a functional recording in which several closely spaced depths are sampled repeatedly. In our continuous-piezo implementation, planes acquired during flyback are excluded from the reconstruction; below, we refer to the remaining planes as the included imaging planes. Unlike z-stack correction, it does not require a separately acquired reference z-stack; instead, a reference volume is constructed from the functional recording itself. The algorithm identifies frames from each imaging cycle that best represent a common target depth and combines them into a depth-stabilized movie, which is subsequently processed with Suite2p for ROI detection and fluorescence extraction (**Figure 4**).

**Figure 4.**
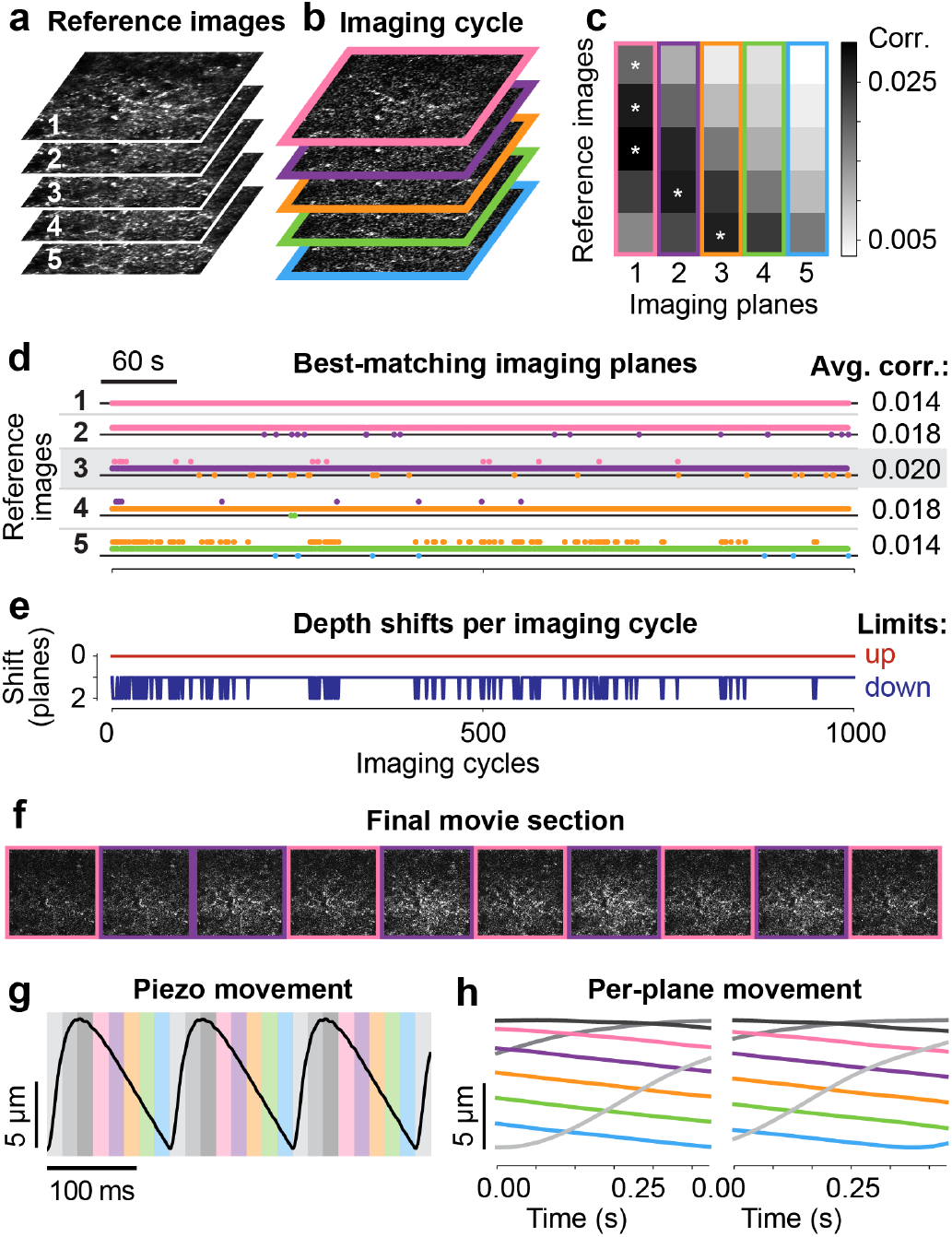
Volume-based z-registration workflow and acquisition geometry. **a**. Reference images for the included imaging planes from an example recording; flyback planes are excluded. **b**. One imaging cycle containing one frame from each included plane. **c**. Correlations between the frames in **b** and reference images in **a**; stars indicate the best-matching frame for each reference plane. The pattern of correlations indicates that the imaging stack sampled in this cycle is shifted downward by two plane spacings relative to the reference volume. **d**. Best-matching imaging plane for each reference plane across the recording; right, mean correlation of the matched frames. Black lines indicate the nominal depth of each reference plane. Across most of the recording, the sampled imaging stacks are shifted downward by approximately one plane relative to the reference volume. **e**. Lower and upper bounds on the axial shift compatible with the frame matches in each imaging cycle. Based on these, reference planes 3–5 remain recoverable throughout the recording. Reference plane 3 has the highest mean correlation and is therefore selected; its depth defines the target depth of the reconstructed movie. **f**. Example frames from the reconstructed depth-stabilized movie; border color indicates the selected source plane. Each output frame incorporates neighboring planes by weighted averaging. **g**. Piezo position during frame recording of several imaging cycles; colored rectangles indicate frame acquisition for included planes and gray rectangles indicate flyback planes. **h**. Mean within-frame piezo trajectories for the imaging planes, illustrating well-aligned (left) and poorly aligned (right) timing between piezo motion and frame acquisition; flyback planes are gray. With poor phase alignment, the lowest included plane (blue) is acquired near the piezo turnaround and has a different within-frame depth trajectory, making it non-parallel to the other included planes.

##### Volume-based z-registration workflow

1. *Build a reference volume from the included imaging planes (**Figure 4**a)* The algorithm constructs one reference image for each included imaging plane from a short block of frames acquired near the temporal midpoint of the recording. Using a temporally compact block limits depth variability during reference construction, while choosing the midpoint places the reference near an intermediate tissue depth when slow drift is approximately monotonic. The resulting per-plane reference images are laterally aligned to form a reference volume.
2. *Match every frame to every reference plane (**Figure 4**b,c)* Each acquired frame is laterally registered to every reference image using the Suite2p registration algorithm, and the correlation is recorded for each frame–reference pair. For each reference plane and imaging cycle, the acquired frame with the highest correlation is identified as its best-matching source frame.
3. *Select a reference plane that remains recoverable across the recording (**Figure 4**d,e)* For each imaging cycle, the algorithm identifies which of the acquired frames best matches each reference plane. These matches are then used to estimate how far the frames imaged during that cycle could have shifted upward or downward in depth, and thus which reference planes are recoverable (see Methods). Across all imaging cycles, the algorithm selects the reference plane that remains recoverable across time. If several planes are equally recoverable, the plane with the highest mean correlation to its matched frames is selected. This reference plane defines the target depth of the reconstructed movie.
4. *Assemble and re-register a depth-stabilized movie (**Figure 4**f)* For each imaging cycle, the algorithm selects the acquired frame that best matches the reference plane chosen as the target depth. This frame is then combined with available frames from the immediately neighboring imaging planes, using weights derived from the similarity of their reference images. This blending reduces discontinuities when the best-matching imaging plane changes between cycles. The assembled movie is then rigidly re-registered in x-y to remove residual lateral offsets and establish a consistent lateral reference.
5. *Detect ROIs and extract fluorescence traces* The final depth-stabilized movie is processed with Suite2p for ROI detection, extraction of ROI and neuropil fluorescence, and optional GUI-based curation. The resulting traces are then processed as for z-stack correction to remove neuropil contamination and calculate ΔF/F (see Methods). No ROI-specific axial correction is required because axial motion has already been corrected at the movie level.

#### Performance and validation

We evaluated volume-based z-registration using multiplane two-photon imaging of retinal ganglion cell axonal boutons in the superior colliculus of awake, head-fixed mice able to run voluntarily on a treadmill. Retinal ganglion cells expressed either synaptically targeted SyGCaMP8m or axon-enriched axon-GCaMP8s. All but one mouse expressed tdTomato in inhibitory neurons, providing a stable, activity-independent signal for lateral and axial registration; in the remaining mouse, GCaMP fluorescence was used instead. A piezoelectric actuator sampled five included imaging planes at 7.5 Hz per plane, with additional planes acquired during flyback and excluded from z-registration (**Figure 4**g,h). Separate recordings were used for ground-truth validation and for quantifying the effects of z-registration on fluorescence measurements.

In two ground-truth recordings, the estimated axial position closely tracked displacements manually imposed with the piezoelectric actuator (**Figure 5**a). The imposed displacements were superimposed on the sawtooth trajectory used for multiplane acquisition. Assuming that brain motion was negligible relative to the imposed displacement, the actuator trajectory provided the ground-truth axial position. During periods in which the selected target depth remained within the sampled volume, RMSEs between estimated and ground-truth positions were 1.03 *µ*m and 0.69 *µ*m in the two recordings. Depth estimates changed in 2-*µ*m increments, corresponding to the spacing between included imaging planes. When the target depth moved slightly outside the sampled volume, estimates remained at the nearest volume boundary, whereas larger excursions produced highly variable estimates. Reliable reconstruction is therefore restricted to periods in which the target depth remains within the sampled volume.

**Figure 5.**
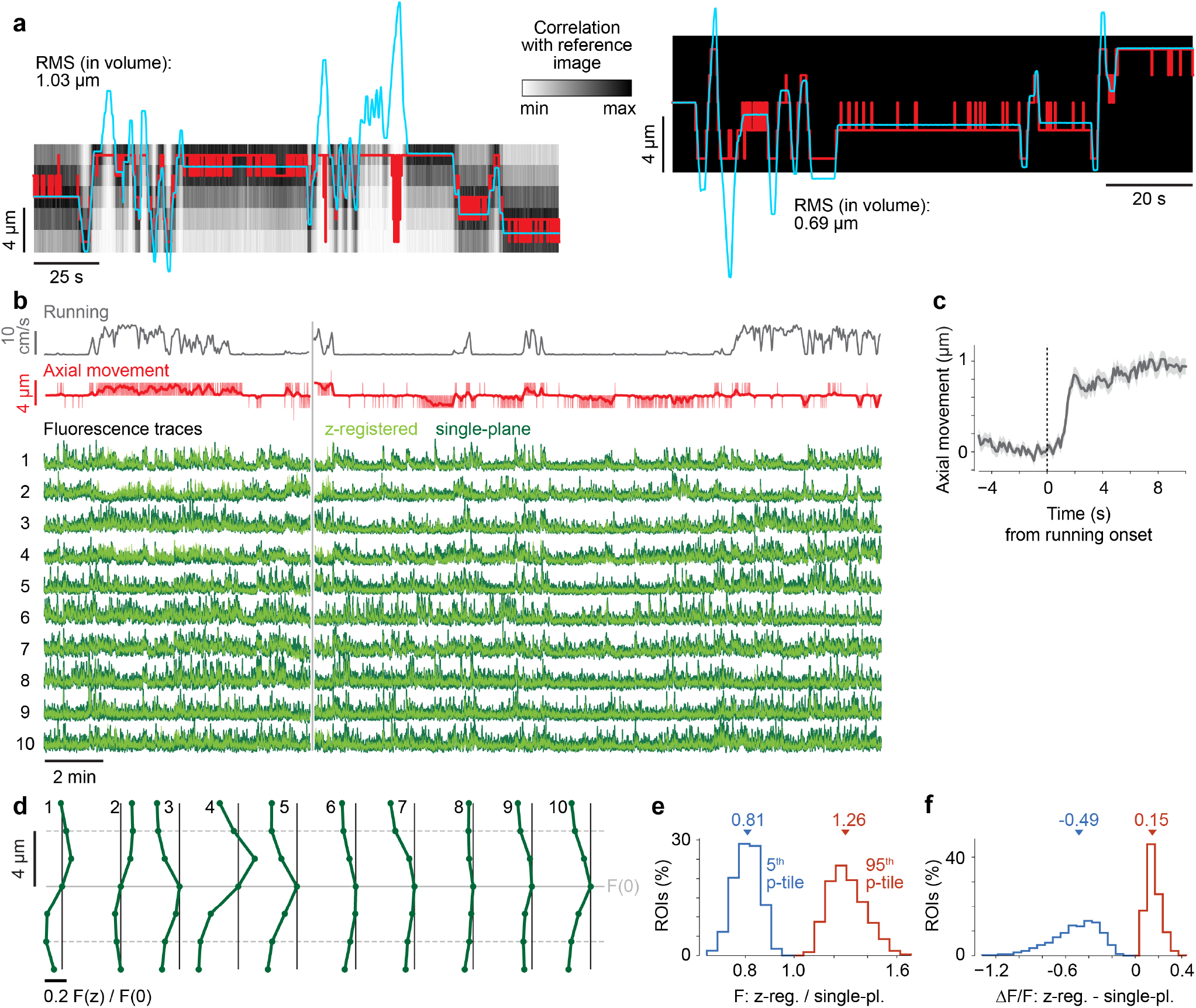
Validation and effects of volume-based z-registration. **a**. Estimated (red) and experimentally imposed ground-truth (cyan) axial displacement in two recordings. Grayscale indicates the correlation of acquired frames with the reference image at the selected target depth. Estimates are reliable only while the target depth remains within the sampled imaging volume. **b**. Running speed, estimated axial displacement, and single-plane (dark green) and z-registered (light green) fluorescence from ten example bouton ROIs. The vertical gray line separates recording blocks using natural images and drifting gratings. **c**. Estimated axial displacement aligned to 80 running onsets from the recording in **b** (mean ± SEM). **d**. Axial fluorescence profiles of the ten ROIs in **b**, showing median single-plane fluorescence at each estimated depth, F(z), normalized to its value at the reference depth, F(0). The solid gray line indicates the target depth and dashed lines the limits of the sampled volume; horizontal scale bar indicates a change of 0.2 in normalized fluorescence. **e**. Across-ROI distributions of the 5th (blue) and 95th (orange) percentiles, calculated over time, of the ratio between z-registered and single-plane fluorescence (2,982 ROIs from six recordings); triangles indicate medians. **f**. Corresponding distributions for the difference between z-registered and single-plane ΔF/F.

In an example functional recording, estimated axial position varied both around running onset and during periods of rest (**Figure 5**b,c). Averaging across 80 running onsets revealed a consistent locomotion-associated axial shift in this recording (**Figure 5**c), similar to the movement observed in the somatic recordings used to evaluate z-stack correction.

The example ROIs exhibited distinct axial fluorescence profiles, illustrating how strongly the fluorescence measured from individual boutons depended on imaging depth (**Figure 5**d). For several ROIs, a displacement of 2 *µ*m from the reference depth changed median fluorescence by approximately 20%, with larger changes at 4 *µ*m.

Across 2,982 ROIs from six functional recordings, we compared fluorescence and ΔF/F extracted from the z-registered movie with traces obtained from the single imaging plane closest to the target depth (**Figure 5**e,f). Using the same summary statistics as for z-stack correction, we calculated the ratio of z-registered to single-plane fluorescence at each time point and summarized its distribution within each ROI by its 5th and 95th percentiles (**Figure 5**e). Across ROIs, the median 5th- and 95th-percentile ratios were 0.81 and 1.26, corresponding to changes of −19% and +26%, respectively. These effects were larger than those observed with z-stack correction in the somatic recordings from the same brain region, consistent with the steeper axial fluorescence profiles of the boutons examined here. We similarly summarized the difference between z-registered and single-plane ΔF/F after neuropil correction and baseline normalization (**Figure 5**f). The median 5th-and 95th-percentile differences were −0.49 and +0.15, respectively.

#### Data requirements and practical considerations

Volume-based z-registration requires a multiplane functional recording in which the same field of view is sampled repeatedly at several closely spaced depths. The included imaging planes must overlap sufficiently in image content, have approximately parallel imaging geometries, and span enough depth that at least one reference depth remains recoverable across the recording.

##### Multiplane acquisition

Images must be acquired at multiple depths in repeated imaging cycles. In our recordings, planes were sampled sequentially while a piezo actuator moved continuously through the imaging volume, and the present implementation was developed and validated for the resulting timing and plane geometry (**Figure 4**g). Other multiplane acquisition approaches may be compatible provided that they repeatedly sample sufficiently overlapping, approximately parallel imaging planes with known plane ordering and geometry.

##### Axial sampling and volume coverage

Neighboring included planes should be spaced closely enough to provide substantial overlap in image content. This allows the algorithm to switch between imaging planes and blend neighboring frames without introducing abrupt changes in the reconstructed movie or extracted fluorescence traces. The appropriate spacing depends on the axial dimensions of the structures of interest and the axial point-spread function of the microscope. For axonal boutons in our recordings, a spacing of approximately 2– 3 *µ*m produced smooth transitions between planes.

The included imaging stack must also span sufficient depth for the target depth to remain represented as the tissue moves axially. If the target depth moves outside the sampled stack, it cannot be reconstructed from the available planes. In our recordings, an axial span of approximately 10 *µ*m was sufficient, although the required coverage will depend on the preparation and expected magnitude of axial motion.

##### Plane geometry and exclusion of flyback planes

Included imaging planes should have approximately parallel geometries, so that switching between planes primarily represents a change in imaging depth rather than a change in the shape or orientation of the sampled plane.

During continuous piezo scanning, plane geometry and spacing depend on the actuator trajectory and its phase relative to frame acquisition (**Figure 4**g,h). The actuator timing should therefore be adjusted so that included planes are acquired during an approximately linear portion of the trajectory, producing similar within-frame depth profiles and approximately constant spacing between successive planes. Frames acquired near the turnaround or during flyback that produce strongly non-parallel imaging planes should be excluded.

The user specifies which imaging planes are excluded. When a piezo-position trace is available, suite2p-in-depth generates a diagnostic plot of position against plane index to help identify flyback planes and assess the geometry of the included planes.

##### Temporal resolution

The reconstructed movie contains one output frame per complete multiplane imaging cycle. Its sampling rate therefore equals the overall frame-acquisition rate divided by the total number of frames acquired per cycle, including flyback frames that are subsequently excluded. Axial coverage therefore comes at the cost of temporal resolution, and sufficiently high frame-acquisition rates are required when measuring rapid calcium dynamics.

The number of acquired planes should be kept no larger than necessary to provide sufficient overlap between neighboring planes and adequate axial coverage. Higher output sampling rates can also be achieved by reducing the number of scan lines, although the field of view may then need to be reduced to maintain adequate spatial sampling.

##### Example acquisition settings

In our recordings of axonal boutons, we acquired eight frames per imaging cycle during continuous piezo scanning, comprising five included imaging planes and three flyback planes. With 256 scan lines per frame and resonant–galvo scanning, the overall frame-acquisition rate was approximately 60 Hz, yielding a final reconstructed-movie sampling rate of approximately 7.5 Hz. The included planes were spaced by approximately 2 *µ*m (**Figure 4**g,h).

#### Assumptions, limitations, and quality control

Reliable z-registration requires accurate frame-to-reference matching and assumes that the sampled imaging stack behaves approximately as a rigid set of planes over each imaging cycle. Additional limitations arise from axial coverage, changes in imaging geometry and the timing of frames within each cycle.

1. Reliable frame-to-reference matching is required Each acquired frame must contain sufficiently stable image structure to permit lateral registration and matching to the reference images. Frame-to-reference similarity should be driven primarily by stationary anatomical features rather than activity-dependent fluorescence, noise or imaging artifacts. Otherwise, the frame with the highest correlation may not be the frame closest in depth to the corresponding reference plane.
2. Estimation of the recoverable depth range assumes limited motion within each imaging cycle To determine which reference planes remain recoverable in each imaging cycle, the algorithm interprets the frame-to-reference matches as arising from a common axial shift of the sampled imaging stack. This assumes that tissue displacement changes little over the duration of a cycle. Substantial movement within a cycle can render the estimated shift range and resulting recoverability bounds unreliable.
3. The reference volume and imaging geometry must remain representative The reference images and geometric relationship between the included planes must remain representative throughout the recording. Changes in the field of view, optical geometry, tissue deformation or actuator trajectory can reduce frame-to-reference correlations or cause different planes to imply inconsistent axial shifts. Such changes cannot be represented by the single axial displacement assumed by the algorithm.
4. Reconstruction is limited to cycles in which the target depth remains within the sampled stack The algorithm can reconstruct a common depth only while the target depth is represented by at least one acquired imaging plane. If axial movement carries the target depth beyond the upper or lower boundary of the sampled stack, that imaging cycle cannot be reconstructed reliably. Frequent best matches to the first or last included imaging plane therefore indicate insufficient axial coverage and limit the usable portion of the recording.
5. Small variations in output-frame timing are ignored The reconstructed movie is treated as uniformly sampled at the nominal imaging-cycle rate. However, when the imaging plane that best matches the target depth changes, the selected frame occurs at a different time within the imaging cycle. These small variations in the interval between successive output frames are ignored in downstream analyses. This approximation is appropriate when the timing differences are small relative to the timescale of the dynamics being measured.
6. Quality-control checks Before interpreting the reconstructed movie and extracted traces, users should inspect frame-to-reference correlations, the imaging plane that best matches the target depth in each cycle, and the frequency with which these matches reach the upper or lower boundary of the sampled stack (**Figure 5**a). The best-matching imaging plane should generally change coherently over time; frequent jumps between non-neighboring planes or inconsistent shifts inferred from different reference planes suggest unreliable depth matching. Persistently low or ambiguous correlations, frequent boundary matches, abrupt plane-transition artifacts or increased noise indicate that z-registration may be unreliable for that recording. suite2p-in-depth automatically generates diagnostic outputs for these checks, including plots similar to those shown in **Figure 4** and **Figure 5**. Guidance for interpreting these outputs is provided in the GitHub repository.

## Discussion

We present two complementary offline approaches for addressing axial motion in functional two-photon imaging, implemented in the Suite2p-compatible package suite2p-in-depth. Z-stack correction uses a reference z-stack to estimate imaging depth and correct depth-dependent fluorescence changes in individual ROIs, whereas volume-based z-registration reconstructs a movie at a common target depth from repeated acquisitions of closely spaced imaging planes. The package also produces diagnostic outputs at the level of the recording and, for z-stack correction, individual ROIs, allowing users to assess depth matching, axial coverage and the effects of correction in their own data.

Both approaches tracked experimentally imposed axial displacements within the sampled depth range. In functional recordings, correction produced modest changes for most somatic ROIs but substantially larger changes in a subset, with larger effects observed in bouton recordings. These results show that the impact of axial motion depends not only on displacement magnitude but also on how the fluorescence of individual structures varies with depth.

### Choosing an axial-motion correction method

The appropriate correction method depends on whether the target structure remains detectable over the expected range of axial motion. Z-stack correction is suitable when the structure remains detectable throughout this range and its axial fluorescence profile provides an unambiguous calibration across depth. It requires a separately acquired reference z-stack but does not reduce the temporal sampling of the functional recording. If axial displacement instead moves a structure far enough that little or none of its fluorescence remains detectable in a single imaging plane, rescaling the measured signal cannot recover the missing information. Volume-based z-registration is designed for this regime: neighboring depths are sampled during each imaging cycle and used to reconstruct a common target depth, at the cost of reduced temporal sampling. Increasing the number of acquired planes extends axial coverage but reduces the output sampling rate at a fixed frame-acquisition rate. Plane spacing, total depth coverage and cycle duration must therefore be chosen according to the axial dimensions of the structures, expected motion and timescale of the signals being measured. Somata and boutons illustrate these different use cases but do not define an absolute boundary between them.

The reliability of correction also depends on the fluorescence available for depth estimation. For z-stack correction, labeling density affects how confidently an axial fluorescence profile can be attributed to a single structure: overlapping structures can produce ambiguous or multi-peaked profiles. When an individual ROI profile cannot provide a reliable depth-dependent calibration, volume-based z-registration may be more appropriate because it maintains a common imaging depth without requiring ROI-specific axial profiles. For both approaches, depth estimation should be based on activity-independent fluorescence whenever possible, because activity-dependent changes could otherwise be mistaken for movement.

Combined axial and lateral registration could potentially be improved by using the prepared reference z-stack directly during Suite2p registration. Each functional frame could first be rigidly matched to its best-corresponding z-stack slice and then nonrigidly registered to that slice, providing a depth-matched reference for lateral registration. This extension is not currently implemented or tested in suite2p-in-depth and would require reprocessing the functional data from the start. Its performance would also depend strongly on the quality and internal alignment of the reference z-stack.

### Assessing the impact of axial motion without correction

Depth estimates and ROI-specific axial fluorescence profiles can also be used to assess the impact of axial motion even when fluorescence correction is unnecessary or cannot be applied reliably. Depth estimates can reveal slow drift, abrupt shifts between recording blocks and periods when a structure approaches or moves beyond its reliably sampled range. Combining these estimates with ROI-specific axial profiles allows the fluorescence error expected from motion to be evaluated relative to the biological effect of interest. A small displacement along a steep axial profile may alter fluorescence substantially, whereas a larger displacement across a relatively flat profile may have little effect.

### Comparison to existing methods

#### Offline approaches

Our two workflows operate offline, leaving the original measurements unchanged. Depth estimates and corrections can therefore be inspected, rejected or repeated without affecting data acquisition.

Related offline methods have likewise used depth-dependent fluorescence profiles to compensate for axial motion (Flores-Valle and Seelig, 2022; Ryan et al., 2020). Ryan et al. (2020) estimated displacement by matching labeled vasculature to a reference z-stack and corrected bouton fluorescence using fitted Moffat profiles. Our method similarly estimates displacement through image matching but uses measured ROI-specific profiles directly and does not require vasculature labeling. This avoids imposing a fixed profile shape, which may be inappropriate for structures with heterogeneous axial profiles. Flores-Valle and Seelig (2022) inferred axial position from two or more simultaneously acquired functional imaging planes. In contrast, our approach does not require simultaneous sampling at multiple functional depths but instead estimates depth relative to a separately acquired reference z-stack. It also supports reslicing of the reference stack to match functional-plane geometry.

Offline registration of repeatedly acquired image stacks has previously been used to reduce motion artifacts during behavior (Andermann et al., 2010), and toolboxes such as CaImAn and Suite3D support three-dimensional registration and volumetric ROI analysis (Giovannucci et al., 2019; Haydaroğlu et al., 2025). Our volume-based z-registration instead reconstructs a single movie at a common target depth and subsequently uses the standard Suite2p workflow for ROI detection and fluorescence extraction. The method builds on a MATLAB extension to Suite2p previously used for retinal bouton imaging (Schröder et al., 2020).

Reconstructing a single target depth is well suited to our shallow, densely sampled recordings, in which neighboring imaging planes contain largely overlapping structures and primarily provide continued access to the same population during axial movement. It also avoids duplicate ROI detections across neighboring planes. By contrast, approaches designed to retain and analyze the full registered volume are more appropriate when distinct populations across depth are themselves of interest.

#### Online acquisition stabilization

Online stabilization provides an alternative by adjusting imaging depth during acquisition to keep structures of interest in focus. This can reduce the additional axial sampling required for offline registration but requires suitable focus-control hardware and tracking software. Tracking errors can also result in irrecoverable signal loss if the structure leaves the sampled region, whereas offline approaches retain the acquired measurements and allow the correction to be inspected or revised.

Chen et al. (2013) used licking-triggered focus adjustments to compensate for axial displacement, relying on a calibrated relationship between behavior and movement rather than direct tissue tracking. Image-based online correction is available in ScanImage, where acquired images are matched to a reference z-stack and estimated axial displacement is compensated by adjusting the imaging depth (Stringer et al., 2026). However, this matching requires compatible sampling geometry between the reference stack and the functional images. During continuous piezo scanning, functional frames can sample oblique planes that differ from those in a separately acquired reference stack. If this geometry is not accounted for, mismatches between functional frames and reference slices can bias depth estimates and the resulting correction. Our offline z-stack correction workflow explicitly accommodates this mismatch by reslicing the reference stack using the recorded piezo trajectory.

Acousto-optic approaches combine rapid three-dimensional scanning with feedback from a fluorescent reference, enabling stabilized recordings from small neuronal structures in behaving animals, including calcium and membrane-potential measurements in awake mouse dendrites (Gonzalez et al., 2026; Griffiths et al., 2020). These systems support rapid, targeted acquisition but require specialized scanning and feedback hardware and a sufficiently bright, stable reference. Our offline workflows can be implemented on standard two-photon microscopes equipped for z-stack or multiplane acquisition and could also complement online stabilization when sufficient spatial information is retained to correct residual axial motion.

#### Optical alternatives

Optical approaches can instead increase tolerance to axial motion by extending the excitation profile in depth, allowing structures to remain visible during displacement without explicitly tracking their position. Bessel-focus imaging, for example, can resolve individual dendritic spines under sufficiently sparse labeling (Lu et al., 2017). With denser labeling, however, integrating fluorescence across depth increases the likelihood that signals from different structures overlap in the projected image. Increased tolerance to axial motion therefore comes at the cost of reduced axial separation of signals. Our workflows retain conventional optical sectioning and instead use additional reference or functional depth samples to estimate and compensate for displacement.

### Conclusion

Together, the presented approaches provide complementary ways to address axial-motion artifacts within a Suite2p workflow. Their applicability is determined not by anatomical identity alone, but by the relationship between the axial extent of the structures, the magnitude of movement and the available depth sampling. By combining depth estimates with recording- and ROI-level diagnostics, *suite2p-in-depth* allows the impact of axial motion to be quantified and correction to be applied only where the available data support it.

## Acknowledgments

We thank Charlotte Donald Wilson for help with mouse husbandry.

This work was supported by a Sir Henry Dale Fellowship jointly funded by the Wellcome Trust and the Royal Society (grant 220169/Z/20/Z to SS) and by the Biotechnology and Biological Sciences Research Council (BB/W014955/1 to SS). For the purpose of open access, the authors have applied a CC BY public copyright license to any Author Accepted Manuscript version arising from this submission.

## Author contributions

Conceptualization: S.S. and L.B.; Investigation: L.B. and M.C.; Software: S.S., A.M., L.B. and C.S.; Data curation: S.S.; Formal analysis: S.S.; Supervision: S.S.; Funding acquisition: S.S.; Writing: S.S. All authors reviewed and approved the final version of the manuscript.

## Data and code availability

The data used to generate the figures in this manuscript are publicly available at doi.org/10.6084/m9.figshare.33981469.

The code for suite2p-in-depth is available at github.com/Schroeder-Lab/suite2p-in-depth.

Scripts for recreating the analyses and figures are available at github.com/Schroeder-Lab/suite2p-in-depth/tree/main/paper.

## Methods

All procedures were conducted in accordance with the Animals (Scientific Procedures) Act 1986 under licenses issued by the UK Home Office following appropriate ethical review.

### Experimental Model and Subject Details

We used 12 mice aged 12–31 weeks: two female Ai14 mice (JAX 007908) and ten Ai14 × Vgat-IRES-Cre mice (six female, four male; JAX 007908 and 016962). The latter expressed tdTomato in inhibitory neurons. Mice were maintained on a 12-h light/12-h dark cycle and were predominantly group-housed.

### Surgical procedures

The surgical preparation, including headplate attachment and implantation of the chronic imaging window over the superior colliculus (SC), was performed largely as described previously (Schröder et al., 2020). Briefly, mice were anesthetized with isoflurane (3.5% induction, 1–2% maintenance). Meloxicam (5 mg/kg), buprenorphine (0.05 mg/kg), and dexamethasone (4 mg/kg) were administered subcutaneously, and bupivacaine (7 mg/kg) was injected beneath the scalp for local analgesia. EMLA cream (5%) was applied to the ear bars and chloramphenicol was used for corneal protection. A 4-mm craniotomy was made over the right SC (approximately −4.2 mm AP and 0.5 mm ML from bregma), and a custom chronic imaging window was implanted as previously described, including anterior displacement of the overlying sinus to expose and stabilize the posterior SC.

For imaging SC neurons, 230 nl of either pGP-AAV2/1-syn-jGCaMP8m-WPRE (Addgene viral prep #162375-AAV1; Zhang et al., 2023) or AAV-hSyn-Soma-jGCaMP8m (Addgene viral prep #169257-AAV1; Grødem et al., 2023) was injected into the posterior right SC at a final titer of approximately 3.5 × 10^12^ GC/ml. Virus was injected 300–500 *µ*m below the brain surface at 2.3 nl every 6 s using a Nanoject II (Drummond), and the pipette was left in place for 5–10 min after injection.

For imaging retinal boutons, 3–4 *µ*l of either AAV2.hSyn.SyGCaMP8m (∼2.5 × 10^12^ GC/ml) or pAAV1-hSynapsin1-axon-jGCaMP8s-P2A-mRuby3 (Addgene viral prep #172921-AAV1; ∼1.5 × 10^13^ GC/ml) was injected intravitreally into the left eye. The SyGCaMP8m construct consisted of GCaMP8m fused to a synaptic-terminal targeting sequence (Dreosti et al., 2009) and packaged in AAV2/2. Injections were performed with a 5-*µ*l Hamilton syringe following pupil dilation with Mydriacil.

Body temperature was maintained at 37°C throughout surgery, and Hartmann’s solution was administered subcutaneously at 0.01 ml/g/h. After surgery, mice recovered on a heating pad and received oral meloxicam for three days. Imaging began at least 3 weeks after SC injection and at least 4 weeks after intravitreal injection.

### Two-photon imaging

Two-photon imaging was performed using a resonant-scanning HyperScope microscope (Scientifica) controlled by ScanImage 2022 and equipped with a 16×, 0.8 NA water-immersion objective (N16XLWD-PF, Nikon). Excitation at 920 nm was provided by a Chameleon Vision II femtosecond laser (Coherent), with approximately 20 mW measured after the objective. Multiplane imaging was performed by continuous piezo scanning with a repeating sawtooth trajectory, using a P-725.4CA PIFOC piezoelectric objective drive (Physik Instrumente). Green GCaMP and red tdTomato fluorescence were detected in separate PMT channels through 525/50-nm and 620/60-nm emission filters, respectively.

For imaging SC neurons, three imaging planes separated by approximately 20 *µ*m were acquired together with one flyback plane. Across recordings, imaging planes were located 30– 185 *µ*m below the SC surface. Fields of view were 360–730 *µ*m across each dimension and were acquired at 512 × 512 pixels and 7.5 Hz per imaging plane.

For recordings used for z-stack correction, a reference z-stack was acquired for each imaging field using the piezoelectric objective drive, either before or after the functional recordings. Z-stacks typically spanned approximately 100 *µ*m in 1-*µ*m steps, with 10 frames acquired at each depth. Field of view, zoom, excitation wavelength and laser power were kept identical to those used for the corresponding functional recording. For depth matching, the tdTomato channel was used when available; otherwise, the GCaMP channel was used.

For imaging retinal boutons, five imaging planes separated by approximately 2 *µ*m were acquired together with three flyback planes. Across recordings, imaging planes were located 15– 90 *µ*m below the SC surface. Fields of view were 140–180 *µ*m across each dimension and were acquired at 256 × 256 pixels and approximately 7.5 Hz per imaging plane.

For ground-truth validation, the piezo offset was manually varied during ongoing multiplane imaging, and the recorded piezo-position signal was used as the imposed axial trajectory.

### Experimental setup

Mice were head-fixed above a 19-cm-diameter treadmill while remaining free to run (Warren et al., 2021). Treadmill rotation was measured with a rotary encoder (1,024 pulses per revolution; Kübler, Germany). Mice were habituated to head fixation for at least seven days before the first recording session, with fixation duration increased gradually from 1 min to 1 h.

During imaging, mice viewed drifting gratings, sparse visual noise, size-varying dots, full-field chirps, natural images, or uniform gray or black screens.

The piezo-position signal and two-photon frame clock were acquired using a PCIe-6323 data-acquisition board (National Instruments), while the rotary-encoder output was acquired through an Arduino Mega 2560 R3 microcontroller. These behavioral and synchronization signals were recorded at 1 kHz. Experimental control and recording of behavioral and synchronization signals were implemented in Bonsai (Lopes et al., 2015).

### Algorithmic details

All processing was implemented in suite2p-in-depth. Unless stated otherwise, image registration and image-similarity calculations used the corresponding Suite2p procedures (Stringer et al., 2026).

#### Z-stack correction

*Reference z-stack construction and alignment* For each z-stack depth, the ten acquired frames were first processed independently of the functional recording. An initial reference image was generated from the five most mutually correlated frames and refined through rigid phase-correlation registration. All ten frames were then rigidly registered to the final reference and averaged. Nonrigid registration was disabled at this stage. When required, bidirectional-scan phase correction was applied before registration.

The resulting averaged z-stack planes were aligned across depth. Each plane was first rigidly registered to the preceding plane, and the resulting pairwise x–y shifts were cumulatively summed and median-centered to minimize the overall displacement of the stack. Following rigid alignment, each interior plane was nonrigidly registered to the average of its two neighboring planes; the first and last planes were registered to their single neighbor. Nonrigid registration used 64 × 128-pixel blocks with a maximum block displacement of 5 pixels.

During continuous piezo scanning, movement of the objective within each frame resulted in functional imaging planes that were not necessarily parallel to the planes of the reference z-stack. To reproduce the geometry of each functional plane, the recorded piezo trajectory was linearly interpolated onto the image rows and the z-stack volume was sampled at the corresponding axial positions. Nearest-neighbor interpolation was used along the z-axis, with positions outside the acquired range clipped to the nearest stack boundary. Each slice of the resulting resliced stack was then registered to the corresponding Suite2p reference image. The rigid and nonrigid x–y transforms obtained for the best-correlating slice were applied identically to the full resliced stack, thereby preserving its internal geometry.

#### Estimation of imaging depth

Each laterally registered functional frame was compared with all slices of the aligned reference z-stack using the same correlation procedure as Suite2p. The depth of the slice with the highest correlation was assigned as the estimated imaging depth for that frame. Matches to either endpoint of the evaluated z range were considered unreliable and set to NaN. The median of the remaining depth estimates across the recording was defined as the reference depth, *z*_ref_.

#### ROI-specific axial fluorescence profiles and correction

For each ROI, the ROI mask was applied to all slices of the aligned reference z-stack to obtain its axial fluorescence profile, *P*_*i*_(*z*). The depth-dependent correction factor for ROI *i* at time *t* was defined relative to the profile amplitude at the reference depth,

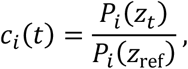

where *z*_*t*_ is the estimated imaging depth. ROI fluorescence and the corresponding neuropil fluorescence were divided by *c*_*i*_(*t*), thereby rescaling both signals to their expected values at the reference depth.

Correction was restricted to a reliable portion of each ROI’s axial profile. Peaks in the normalized profile were detected using `scipy.signal.find_peaks` with `distance=7, width=2, wlen=31`, and `rel_height=0.5`. The minima between neighboring peaks, or between a peak and the corresponding stack boundary, were used to delimit the profile region surrounding the reference depth. If no peaks were detected, no peak-based restriction was applied. Independently of these boundaries, depths at which the normalized axial-profile amplitude fell below 0.5 were considered unreliable. Fluorescence measurements for which the estimated depth fell outside the resulting reliable region were set to NaN rather than extrapolated.

#### Neuropil correction and ΔF/F

After axial correction, frames marked as invalid were set to NaN and the configured PMT dark offset was subtracted from the ROI and neuropil fluorescence traces. Let *F*(*t*) and *N*(*t*) denote the resulting ROI and neuropil fluorescence, respectively. Slow baselines, *B*_*F*_(*t*) and *B*_*N*_(*t*), were estimated separately for each experimental block. With the settings used here, the 8^th^ percentile was calculated in a centered window of approximately 30 s and the resulting trace was smoothed with a one-dimensional Gaussian with a standard deviation of 60 s. Baseline estimation did not cross boundaries between experimental blocks. Residual fluorescence was then calculated as

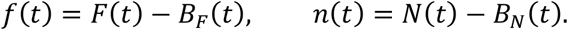

To estimate neuropil contamination for each ROI, the range between the 10th and 90th percentiles of *n*(*t*) was divided into 20 equal-width bins. For each populated bin, the 5^th^ percentile of *f*(*t*) was calculated. These values were regressed against the corresponding neuropil-bin lower edges using ordinary least-squares linear regression,

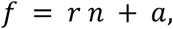

where the fitted slope *r* was restricted to the interval 0–2 and the intercept *a* was unrestricted. Neuropil-corrected fluorescence was calculated as

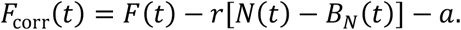

The ROI baseline calculated before neuropil correction was retained for calculation of ΔF/F:

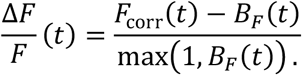

Thus, no new baseline was estimated after neuropil correction; instead, the denominator was bounded below at one fluorescence-intensity unit to avoid numerical instability.

### Volume-based z-registration

#### Reference-volume construction and frame matching

Planes acquired during piezo flyback were excluded before z-registration. A reference image was generated for each remaining imaging plane using the standard Suite2p reference-image procedure. The number of contributing frames was user-specified as in Suite2p (default 400), except that consecutive frames near the temporal midpoint of the recording were used rather than frames sampled across the recording. The resulting per-plane reference images were laterally aligned to form a reference volume.

Each acquired frame was then rigidly registered to every reference-plane image using Suite2p registration, and the corresponding correlation was stored. Let *C*_*t,r,p*_ denote the correlation in imaging cycle *t* between candidate reference plane *r* and the frame acquired from plane *p*. For each imaging cycle and reference plane, the best-matching acquired plane was

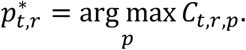

##### Selection of the target reference plane

The displacement implied by each reference-plane match was expressed in units of included imaging planes as

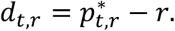

For each imaging cycle, lower and upper bounds on the inferred axial displacement were defined as

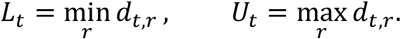

These bounds represent the range of axial shifts compatible with the observed best matches across reference planes.

For candidate target plane *k*, recoverability was scored from whether the two displacement bounds remained within the *P* included imaging planes:

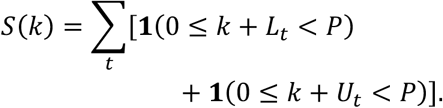

Thus, the two endpoints contributed independently, giving each imaging cycle a score of 0, 1, or 2 for each candidate plane. The plane with the highest total endpoint-coverage score was selected as the target reference plane. If several planes had the same score, the plane with the largest mean best-match correlation,

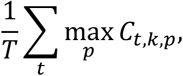

was selected. All calculations were performed in plane-index units rather than physical z coordinates.

##### Reconstruction of the depth-stabilized movie

To reduce discontinuities when the best-matching source plane changed, each reconstructed frame incorporated the selected imaging plane and, where available, its two immediate neighbors. Let *b* denote the selected global reference plane. Each reference image was phase-correlated with reference image *b*, yielding correlation scores *c*_*i*_ Static weights for *b* − 1, *b*, and *b* + 1 were calculated as

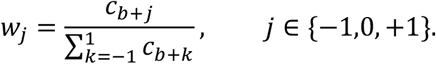

For imaging cycle *t*, let *s*_*t*_ denote the imaging plane that best matches the target plane *b*, and let *X*_*i,t*_ denote the registered frame acquired from plane *i* in that cycle. The reconstructed frame was then obtained from the registered source frames surrounding *s*_*t*_:

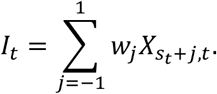

The weights were determined once from the reference-image similarities and remained fixed over time. If a neighboring plane was unavailable or its frame was marked as bad, its weight was set to zero and the remaining weights were renormalized before reconstruction.

Finally, the reconstructed movie was rigidly registered in x–y using the standard Suite2p rigid-registration procedure without additional non-default settings. The resulting movie contained one frame per complete imaging cycle and was subsequently processed with Suite2p for ROI detection and fluorescence extraction. Neuropil correction and ΔF/F calculation were performed as described above.

## Notes

https://doi.org/10.6084/m9.figshare.33981469

https://github.com/Schroeder-Lab/suite2p-in-depth

### Competing Interest Statement

The authors have declared no competing interest.

